# Mass Dynamics 1.0: A streamlined, web-based environment for analyzing, sharing and integrating Label-Free Data

**DOI:** 10.1101/2021.03.03.433806

**Authors:** Joseph Bloom, Aaron Triantafyllidis, Paula Burton (Ngov), Giuseppe Infusini, Andrew Webb

## Abstract

Label Free Quantification (LFQ) of shotgun proteomics data is a popular and robust method for the characterization of relative protein abundance between samples. Many analytical pipelines exist for the automation of this analysis and some tools exist for the subsequent representation and inspection of the results of these pipelines. Mass Dynamics 1.0 (MD 1.0) is a web based analysis environment that can analyze and visualize LFQ data produced by software such as Maxquant. Unlike other tools, MD 1.0 utilizes cloud-based architecture to enable researchers to store their data, enabling researchers to not only automatically process and visualize their LFQ data but annotate and share their findings with collaborators and, if chosen, to easily publish results to the community. With a view toward increased reproducibility and standardisation in proteomics data analysis and streamlining collaboration between researchers, MD 1.0 requires minimal parameter choices and automatically generates quality control reports to verify experiment integrity. Here, we demonstrate that MD 1.0 provides reliable results for protein expression quantification, emulating Perseus on benchmark datasets over a wide dynamic range.

The MD 1.0 platform is available globally via: https://app.massdynamics.com/.

## Introduction

Proteomics can be defined as the application of technologies for identification and quantification of protein content of complex biological samples. Over the past few decades, proteomics research and the complexity of experimental research questions addressed by it have developed rapidly and are having a growing impact in biological and medical research. In particular, areas such as understanding mechanisms of action in disease progression and therapeutic intervention, as well as detection of diagnostic markers, identifying candidates for vaccine production, understanding pathogenic mechanisms and gene expression patterns have been of growing importance in advancing many areas of medically related research^1,2^.

Mass Spectrometry via LC/MS-MS is the leading technology in proteomics research, with Label Free Quantification (LFQ) and isotope-based labelling methods being two approaches that facilitate interrogation and measurement of a large number of analytes in even very complex samples. Isotope based labelling methods such as SILAC and TMT have provided the gold-standard for protein quantification, but are limited by their applicability to types of samples which cannot be easily labelled and by the associated cost of reagents required. By contrast, LFQ is a simpler, more economical and scalable method that requires considered experimental design in order to achieve robust biological insights^3^.

An exemplar of the growing use and adoption of LFQ based approaches is the sheer variety of analytical software packages that have been developed to support LFQ experiments which have been comprehensively covered recently by Tang et. al.^4^, the most widespread of which is MaxQuant^5^. MaxQuant’s success can be attributed at least in part to the holistic nature of it’s analytical tools, covering a breadth of steps including feature extraction, database search, protein identification and quantification which must otherwise be achieved using a combination of other tools.

While data processing is essential for the success of proteomic analysis, objective quality control reporting and reproducible downstream analysis is equally important. The Perseus computational platform is the analytical counterpart to MaxQuant and provides users with a highly flexible framework for post-processing and visualising results^6, 7^. Despite the diverse suite of tools offered by Perseus (accompanied by supporting documentation and online tutorials), the sheer amount and breadth of functionality can be overwhelming for users requiring straight forward analysis of LFQ proteomics data. For some users, who commonly repeat identical analytical procedures frequently for collaborators, Perseus can be more labour and time intensive than other tools designed to automate the same process such as LFQAnalyst ^6,8^.

Since the publication of LFQAnalyst in 2019, there have been numerous publications of automated LFQ pipelines including Eatomics ^9^ and ProVision ^9,10^. While each of these resources differ in their scope and degree of automation, they are among a growing list of attempts to shift toward standardization through automationstraightfoward statistical analysis and visualization performed as an output of the LFQ pipeline. Whilst these attempts have undoubtedly made headway toward more accessible and reliable analytical processes, there still remain large challenges such as scale, storage, sharing and platform sustainability that will inhibit broad adoption among the community. MD 1.0 addresses these needs by meeting multidimensional criteria in reliability, ease-of-use and transparency while offering functionality designed to facilitate collaboration and sharing. A qualitative survey of the tools mentioned above and MD 1.0 is provided in supplementary **table S1**.

There are a number of measures that can be taken to ensure that tools facilitating LFQ proteomics maintain reliability of analysis and interpretability of results. Firstly, data analysis and statistical approaches should be demonstrated to produce accurate and reliable results in accordance with accepted best practice of statistical testing. Secondly, by demonstrating that the specific implementation of these methods has been successful and continues to be over the life of the tool. This ensures that the specific code works and is not accidentally compromised during further development. Thirdly, by incorporating automated quality control metrics and figures. These provide confidence that results on non-benchmark datasets can be reasonably interpreted. By meeting these three criteria, any scientific tool can be safely shared with users of varying degrees of expertise.

However, reliability of results are only one side of the coin when it comes to developing useful, empowering tools for the proteomics field. Interviews with 100 Scientists in the field of Mass Spectrometry^11^ found that “free” tools often imposed significant hidden costs in time spent learning to use the software and in manual tuning of parameters. Though opportunity costs of users are hard to quantify, they represent a genuine cost of use.

It is likely that an awareness of these hidden costs is in part a large driver for the number of tools created for automating analysis of LFQ data, although calls for robust, user-friendly automation in proteomics have been present since 1999^12^. Automated tools are not only less time consuming but reduce the number of points of failure that must be investigated in development, benchmarking and comparison. For many practical reasons, automation simplifies tools, making them more accessible to users and scientific developers.

Moreover, codifying analysis and open sourcing analysis and benchmarking allows for standardisation that addresses reproducibility difficulties that plague the broader scientific field.

Tools which automate highly complicated tasks inevitably end up comprising large amounts of code that must be available for the tool to be reproducible (as a necessary, but not sufficient condition). Tools such as OpenMS^13^ achieve reproducibility by open sourcing their code enabling thorough and ongoing peer review, as tools grow over time.

MD 1.0 attempts to meet these standards of reliability, automation (thereby ease-of-use) and transparency in the specific domain of LFQ analysis. However, MD 1.0 provides further functionality designed to assist with resource integration, annotation, sharing and collaboration.

Traditional and ubiquitous methods for sharing LFQ data likely include email or messenger service (eg: Slack) type transfers of text and use of file storage tools like Dropbox or Google Drive. These tools present obstacles to effective collaboration such as separating data from quality control metrics which can obscure interpretability. By allowing an entire experiment to be shared with ease, MD 1.0 attempts to make supervision and collaboration easier for scientists to ask for and receive assistance while performing their experiments.

## Methods

### Raw data processing with MaxQuant

MaxQuant v1.6.17.0 was used with default parameters except for an LFQ min. ratio of 1 and enabling match between runs. Output files used for quality control include msms.txt, peptides.txt, modificationSpecificPeptides.txt, proteinGroups.txt and evidence.txt whilst only proteinGroups.txt was used for quantification. Where maxquant labelled spiked proteins as potential contaminants, proteinGroups.txt was manually edited to remove this label so these rows would be present in subsequent processing.

The results and parameters files can be downloaded from the supplementary files and from the the MassDynamics platform at the following addresses:

iPRG2015: https://app.massdynamics.com/p/56a6d7f7-c129-4d7a-a221-d2ef3b8ff4c3
Dynamic Benchmark Range Dataset: https://app.massdynamics.com/p/152bd07f-ddd6-4943-bf62-298648b43bd3
HER2: https://app.massdynamics.com/p/b832f17a-a9f3-4890-8533-2d69835f814e

Please note that public experiment links provide a limited feature set due to “read only access”. Downloading input files from the supplementary information and uploading as new experiments will allow access to all features.

### MD 1.0 *LFQ Processing*

Statistical analysis is performed using R version 3.6.0.

An experiment design file is generated from user input during experimental set up in the application prior to processing which is thereafter automatic.

Proteins corresponding to reverse sequences, potential contaminants and proteins only identified by site are filtered out. Intensities provided inside proteinGroups.txt are converted to log2 scale and values imputed using the MNAR (“Missing Not at Random”) method with a mean shift of negative 1.8 and a standard deviation of 0.3 as recommended by in the Perseus protocol. The probability of differential expression was calculated using the function *lmFit* from the Bioconductor package *limma* followed by *eBayes* using the default settings with the exception of “robust” and “trend” that are set to TRUE and false-discovery rate correction using the Benjamini–Hochberg method Protein groups where more than 50% of intensities are imputed for both conditions are excluded from the quantitative analysis. P-values are adjusted using the Benjamini Hochberg approach.

This code is provided in the following github repository: https://github.com/MassDynamics/lfq_processing

In application graphics are produced indicating that an FDR cutoff of 5% and fold change of at least 2 are required for a result to be significant.

### Implementation

MD 1.0 is composed of two separate components, a web component using a modern software stack (javascript and rails) and a processing component built using R and Elastic Compute Cloud (ec2) on Amazon Web Services (AWS). The combination of these two components ensures that processing is repeatable and runs in a computing environment using identical software, parameters and code, while maintaining the privacy & security of user data. Users have the option to share their data with specific users or share publicly.

### Perseus Processing

All proteinGroups.txt were analysed with Perseus version 1.6.14.0. Proteins corresponding to reverse sequences, contaminants and proteins only identified by site were removed. Intensities were transformed to the Log2 scale and missing values imputed as in the MD 1.0 protocol. Student t-tests were performed for each pairwise comparison with an S value of 0 and a Benjamini Hochberg FDR correction. Tests were only performed when more than 50% of values aren’t imputed in at least one group. Session files are available in the supplementary material.

### Benchmarking Datasets

Different LFQ benchmarking datasets are chosen to verify that MD 1.0 recovers near identical results to Perseus including two datasets with ground truth PXD000279^3^, PXD002057^14^ and one without, which constitutes a “real world” scenario PXD010981^15^.

PXD000279 (“dynamic range dataset”) contains raw data for 2 samples (4 replicates each) enriched with one of two “Universal Protein Standards” (1 and 2) which test LFQ accuracy over a large dynamic range.

PXD002057 (“HER2 dataset”) contains raw data for an experiment with 2 samples, each with 3 replicates. These two samples come from 2 cancer cell lines, a parental SKBR3 cell line and another cell line derived from the first which is resistant to human epidermal growth factor receptor 2 (HER2)-targeted therapy.

Lastly, PXD010981 (“iPRG2015 dataset”) contains raw data for the iPRG2015 benchmarking dataset. This dataset is composed of 4 samples with 200ng of tryptic digests of S. cerevisiae (ATCC strain 204508/S288c) were then spiked with different quantities of six individual protein digests of Ovalbumin, Myoglobin, Phosphorylase b, Beta-Galactosidase, Bovine Serum Albumin and Carbonic Anhydrase according to a schema in that publication (present in the benchmarking github repository).

In order to ensure that certain spiked proteins (P02768, P06396 for the dynamic range dataset data and P44683 for the iPRG2015 data) were not removed as contaminants, proteinGroups.txt output were manually edited prior to benchmarking analysis. The resulting input files are provided in the supplementary information.

### Figure Generation and Results Comparison

All results figures and tables were calculated using bespoke python scripts utilizing packages including pandas for data manipulation, plotly and matplotlib for graphics and scipy for pearson correlation calculations.

This code is provided in the following github repository: https://github.com/MassDynamics/lfq_benchmark

## Results and Discussion

### Overview of MD+ Discovery, User Interface, Experiment Creation and Sharing

Mass Dynamics 1.0 is a web-based, integrated and automated analysis and collaboration environment that facilitates LFQ experiments. MD 1.0 enables users to upload MaxQuant output files and visualize the quantitative analysis.The user interface begins at the Experiments page, after users sign up and log in (**Figure 1)**.

**Figure 1.**
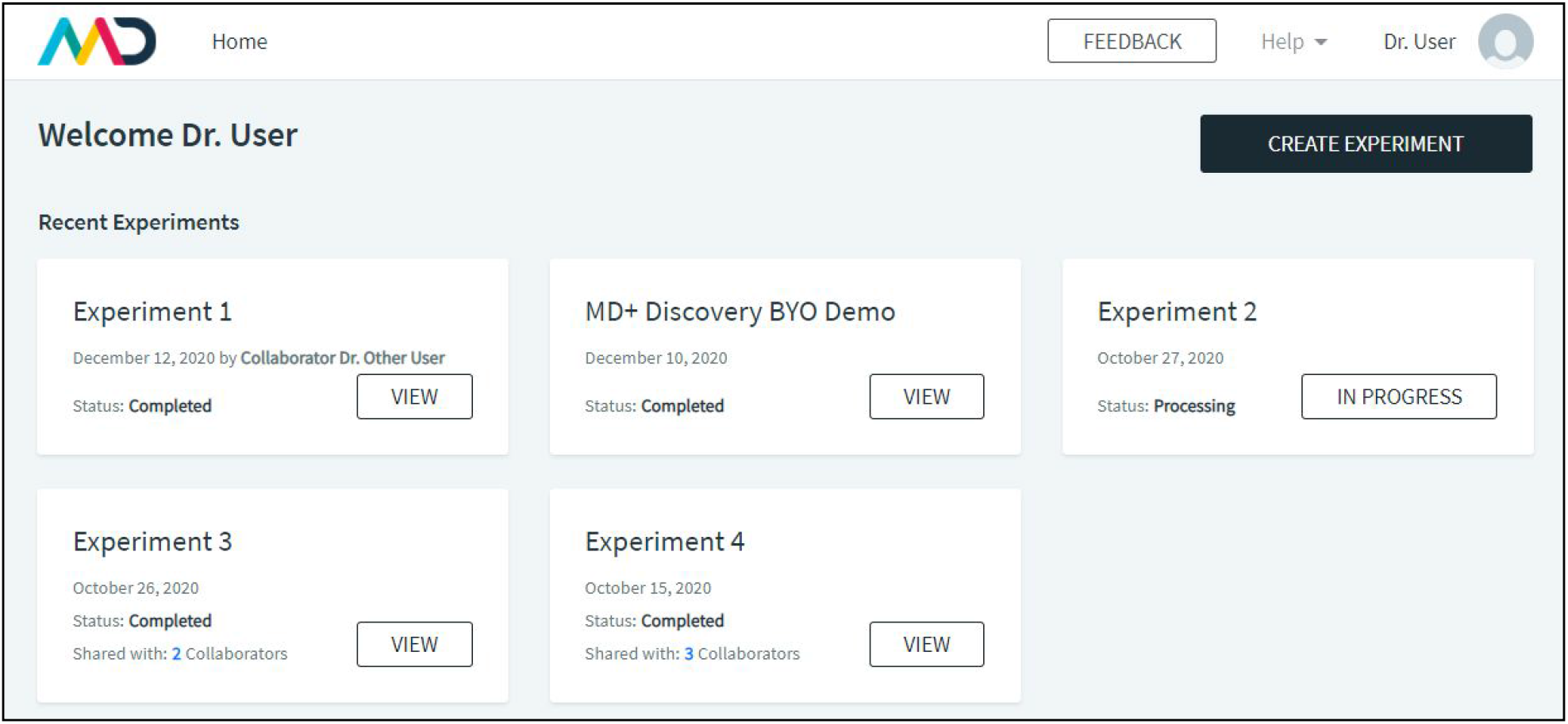
MD 1.0 Landing page. The user interface begins at the experiments page, after users sign up and log in. Users can see their experiments, with associated status (in progress or “view” for finished experiments), dates and owners. Experiments be the users own (such as Experiments 2) or shared via a collaborator (Experiment 1). Users can see which of their experiments have been shared with other users, and who those users are via the “Shared with” link in bold. Experiments are either “in progress” or completed in which case the “view” button is accessible. The “create experiment” button allows users to upload data for a new LFQ analysis.

To create a new experiment, users can click the Create Experiment button on the landing page which takes them to the experiment creation page. Here they are prompted to upload their MaxQuant .txt files which will be used for quantification and quality control reporting. After this, experiment files are presented to the user, who is prompted to allocate them into replicate groups. They can then add an experiment description and complete the experiment creation step. Computing usually takes a matter of minutes, with users being sent an email notification when their experiment has been completed.

In an experiment view (**Figure 2**), users can choose between the following tabs: Analysis, QC Report, Insights, Results Files, Export and Experiment Design. Analysis contains protein expression volcano plot and tables. The QC Report contains a number of experimental quality control plots. The Insights tab contains a list of comments and notes introduced by the user. The Results Files tab contains a list of output files that can each be downloaded, which includes the tab separated files produced by the quantitative analysis scripts. The Export tab enables users to export the volcano plot graph with all “candidate proteins” (proteins that have been selected and added to at least one selection list) highlighted and annotated. Lastly, the Experimental Design tab indicates which files have been assigned to each experimental condition.

**Figure 2.**
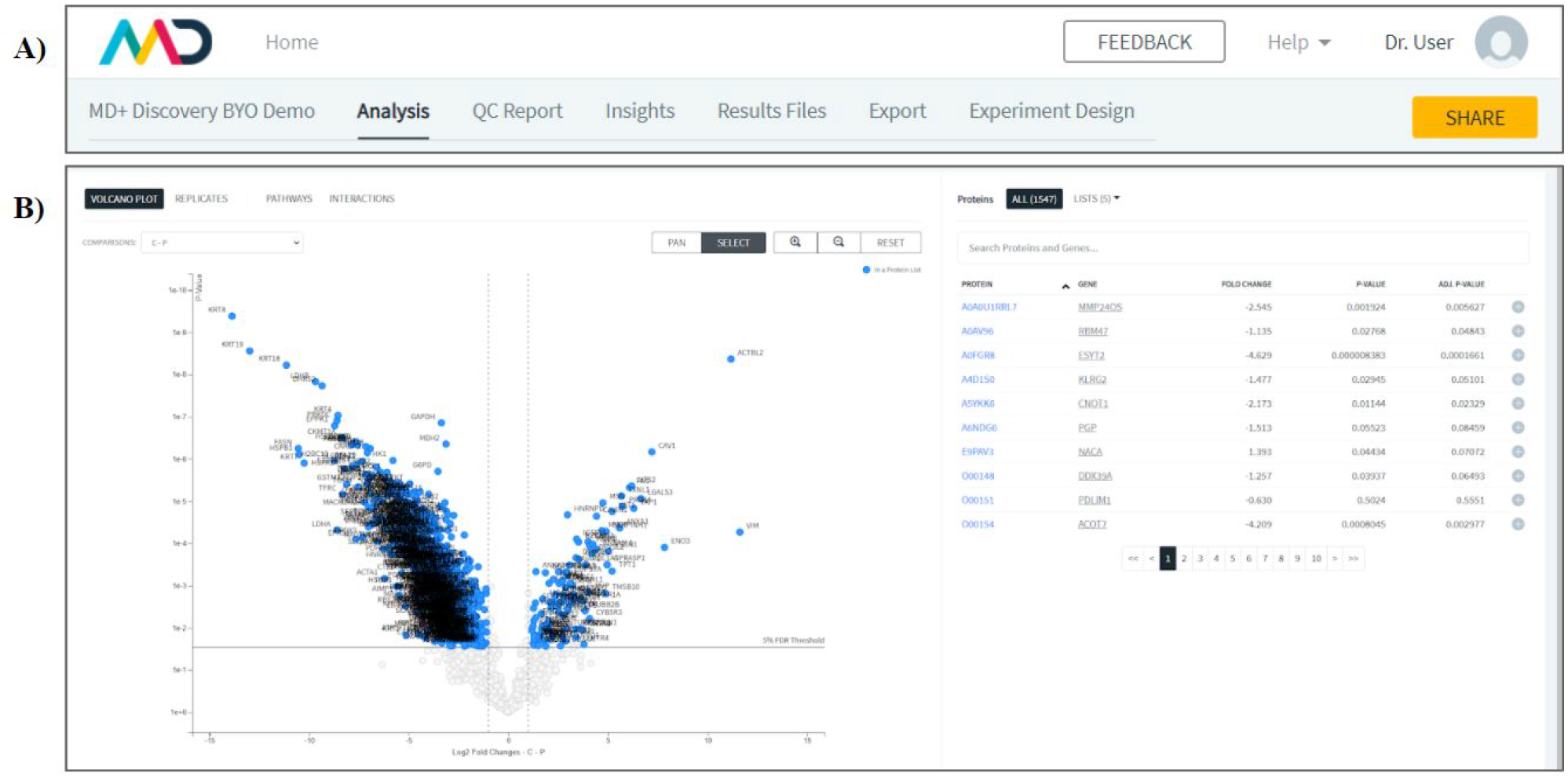
MD+ Discovery Experiment View header (2a) and Experiment View Volcano Plot (2b). In an experiment view (2a), users can choose between the following tabs: Analysis, QC Report, Insights, Results Files, Export and Experiment Design. Users also have the ability to share their experiment. Inside the analysis tab (2b), users can choose between tabs for viewing the volcano plots, filtering by protein observations by replicates and the pathways tab. The volcano plot and table allows users to dynamically, search for, select and manipulate proteins lists by adding, removing and annotating proteins as they complete their analysis.

As the experiment data is stored securely in the cloud, sharing is as simple as giving collaborators access to the same web site. After an experiment is completed, users are able to select the Share button and enter the email address of someone they would like to share their experiment with. This email address is then sent an invitation to join MD 1.0 which contains a link to the shared experiment. Insights associated with particular proteins, which are accessible at all protein level analysis interfaces, perpetuate between users accessing the same experiment and are thus shared with the experiment data.

### MD 1.0 *facilitates analysis and annotation of Label Free Quantitation Results*

After an experiment has finished processing, the Analysis tab can be used to visualize results in the form of a volcano plot (**Figure 2b**). Next to the volcano plot is a table containing a list of each protein with it’s gene, estimated fold change, p-value and adjusted p-value. From this table, proteins can be added to lists which can be used to filter results to proteins of interest or into groups of up or down regulated proteins. A protein in a list can be given an annotation, where the user is able to comment on a protein with text and/or hyperlinks which are then presented in the Insights tab.

### MD 1.0 *allows users to perform Over-Representation Analysis with Reactome*

While MD 1.0 automatically links protein accession codes from table views to Uniprot.org^16^, the Pathways tab (**Figure 3**) provides further integration with an external knowledge base. The Pathways tab uses the Reactome^17^ API content service to provide Over-Enrichment Analysis (ORA) results to users. For each candidate list, one API call is made to perform ORA, while a second is used to retrieve the complete list of proteins in each resulting pathway. A Volcano plot is shown with all proteins in that pathway and that experiment and the “candidate proteins” labelled, with corresponding estimated False Discovery Rate (FDR) provided.

**Figure 3:**
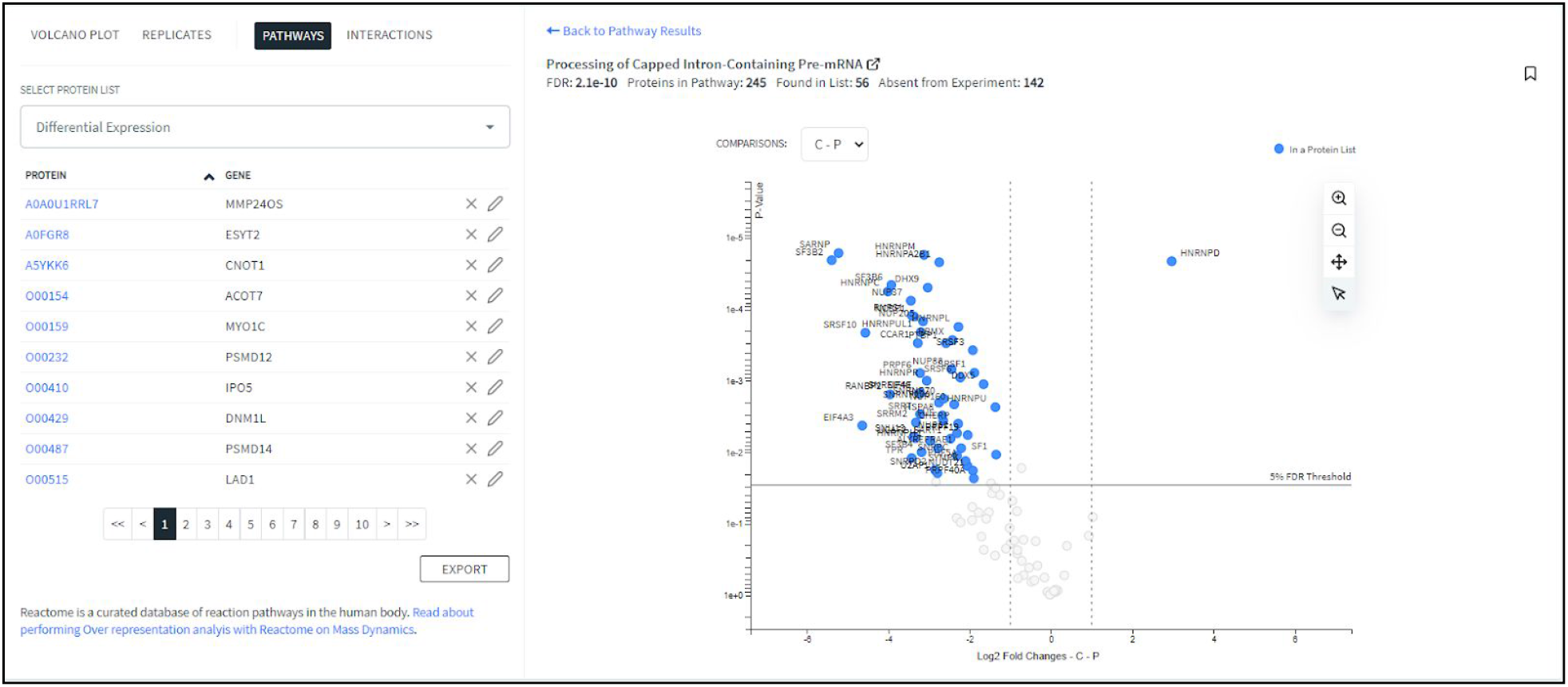
The Pathways view enables users to perform over-representation analysis (ORA) using the Reactome API. As no background can be specified using the Reactome API, the hypergeom test is performed using the ratio calculated as the number of entities in the pathway and in the candidate list, divided by the total number of associated entities known to Reactome. The provided FDR is calculated using the Benjamini-Hochberg method.

### MD 1.0 *Quality Control Report produces diagnostic figures to assess experiment health*

A feature of MD 1.0 is the automated generation of a quality control report accessible in the Quality Control (QC) Report (**see Supplementary Files**) tab in the experiment view.

The report contains three sections, Experiment Health, Feature Completeness and Identifications. Experiment Health contains principal component analysis scatter plots of the first two principal components and a scree plot for all proteins, differentially expressed proteins, modification specific peptides and all peptides. Quantitative CV (coefficient of variation) distributions and Sample Intensity Correlation plots are then produced for proteins, peptides and modification specific peptide tables. The Feature Completeness section provides the percentage of missing measurements in a histogram at the protein, peptide and modification specific peptide levels and a histogram of the percentage of all measurements missing at the LC-MS run (file) level. Lastly, the Identifications section reports counts per file for detected PSMs, modification specific peptides, peptides and proteins. Complete QC reports are provided both in application and in the supplementary information.

Due to the automatic nature of the QC report and data processing, users are able to quickly review and assess experimental results and confidently interpret results or share with collaborators via other features.

### MD 1.0 *reproduces Perseus Results Reliably on Sample Datasets*

In order to determine the reliability of the MD 1.0 automated workflow for LFQ quantification, we analysed the same experiments on Perseus and MD 1.0. Two of these experiments, the iPRG2015, and dynamic range dataset datasets contained ground truth data whereas the last dataset from a study on breast cancer resistant cells constitutes a more realistic real world scenario with no ground truth.

We compared Perseus and MD 1.0 results on these datasets in continuous and discrete measures of accuracy (for the ground truth datasets) and in similarity (for all datasets).

**Table 1** and **Table 2** show the confusion matrices for Perseus and MD 1.0 performance across iPRG2015 and dynamic range dataset data sets respectively. We defined a “true positive” if the estimated and true protein abundance ratios were greater than 2 in the same direction and an adjusted p-value of less than 0.05 was produced. “False negatives”, where the true protein abundance ratio was greater 2 but the estimated value was not or the adjusted p-value was greater than 0.05, were rare but occured once in the dynamic range dataset (in common between the two experiments) and three time for perseus in the iPRG2015 dataset. **Table 1** and **Table 2** provides more details on these missed detections.

**Table 1:** Binary Evaluation of Differential Expression Predictions between Persus and MD 1.0 on Dynamic Range Benchmark Dataset (Cox et al.) Confusion matrices for the dynamic range dataset dataset were identical between MD 1.0 and Perseus. Both methods produced results within the 1% standard for an acceptable false discovery rate. One false negative was produced by both Perseus and MD 1.0 pertaining to Gamma-synuclein (UniProt ID: O76070), resulting from higher adjusted p-values of 0.578975 and 0.351076 respectively.

|  | MD 1.0 |  | Perseus |  |
| --- | --- | --- | --- | --- |
|  | expected true | expected false | expected true | expected false |
| observed true | 39 | 4 | 39 | 4 |
| observed false | 1 | 2198 | 1 | 2198 |
| sum | 40 | 2202 | 40 | 2202 |

**Table 2:** Binary Evaluation of Differential Expression Predictions between Persus and MD 1.0 on iPRG2015 Benchmarking dataset (et al.) Confusion matrices for the iPRG2015 results. Both methods produced results within the 1% standard for an acceptable false discovery rate. The Perseus protocol used failed to detect 3 protein abundance ratios greater than 2, in all cases due to lower confidence than required by the definitions used. The comparisons were: Phosphorylase b in samples 1 vs 2 (adjusted p-value 0.058), Ovalbumin in samples 2 vs 3 and 3 vs 4 (adjusted p-value 0.0594 and 0.0880).

|  | MD 1.0 |  | Perseus |  |
| --- | --- | --- | --- | --- |
|  | expected true | expected false | expected true | expected false |
| observed true | 30 | 14 | 27 | 3 |
| observed false | 0 | 19528 | 3 | 19539 |
| sum | 30 | 19542 | 30 | 19542 |

Pearson correlation was used to measure the similarity between estimated log fold change and the true log fold change (according to the benchmark dataset descriptions). Scatter plots of these values are provided in (**Figure 4a-d**). In all cases pearson correlation was greater than 0.9 suggesting very high accuracy.

**Figure 4.**
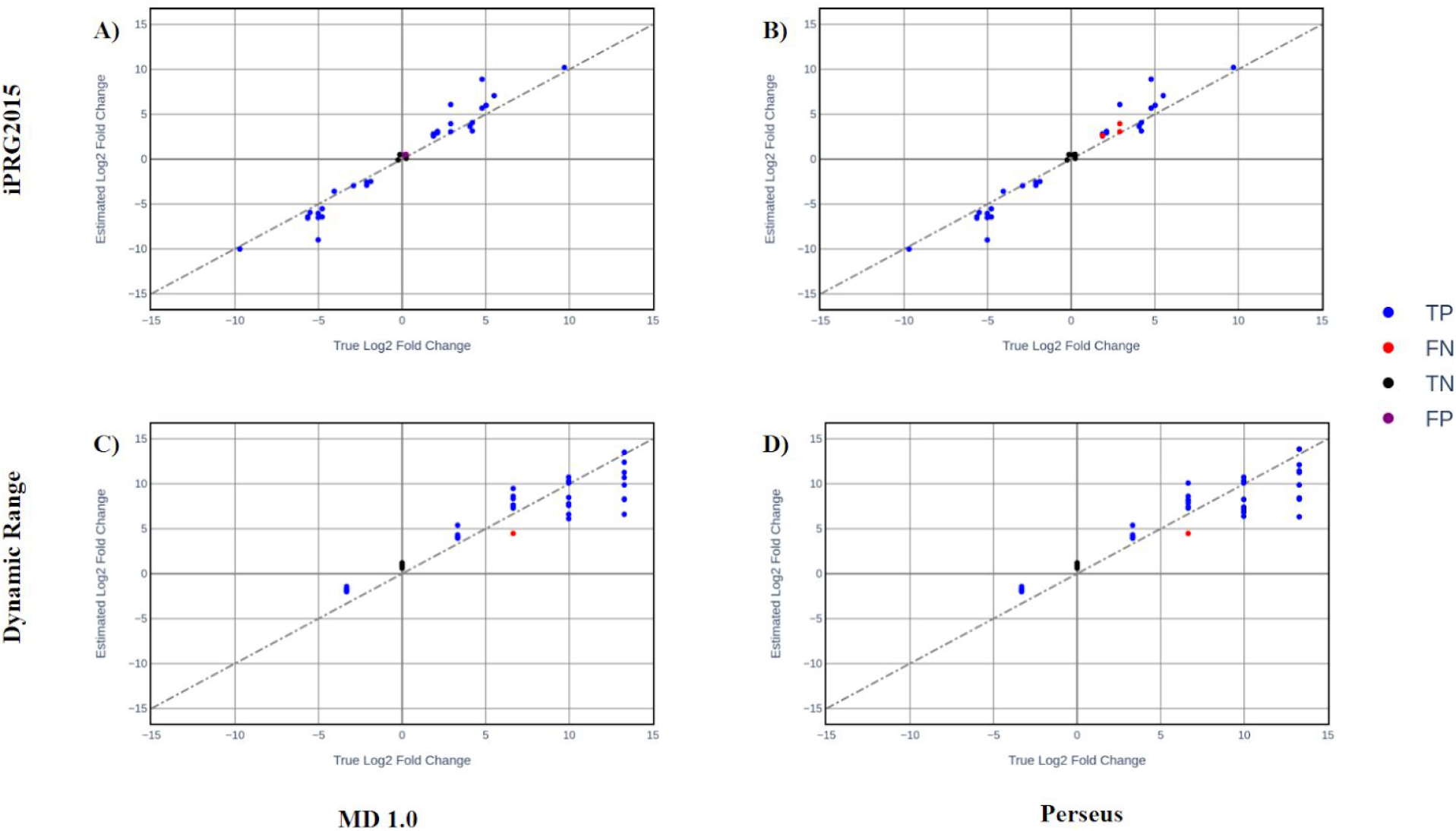
Scatter plots of the true and estimated log fold changes produced by Perseus, MD 1.0 for iPRG2015 and the dynamic range dataset datasets. (A) iPRG2015 Perseus (Pearson correlation = 0.978) (B) iPRG2015 MD 1.0 (Pearson correlation = 0.978) (C) dynamic range dataset MD 1.0 (Pearson correlation = 0.942) (D) dynamic range dataset Perseus (Pearson correlation = 0.938).

The pearson correlation between MD 1.0 and Perseus estimated log fold changes were 0.998, 1.0 and 0.901 for the dynamic range dataset, iPRG2015 and HER2 studies respectively (**Figures 4 - 6**) whereas the associated log10 adjusted p-values varied slightly more, with pearson correlations 0.925, 0.978 and 0.901 for the dynamic range dataset, iPRG2015 and HER2 studies respectively, showing that MD 1.0 produces results consistent with what can be achieved by those using the Perseus platform except with a completely automated workflow.

**Figure 5.**
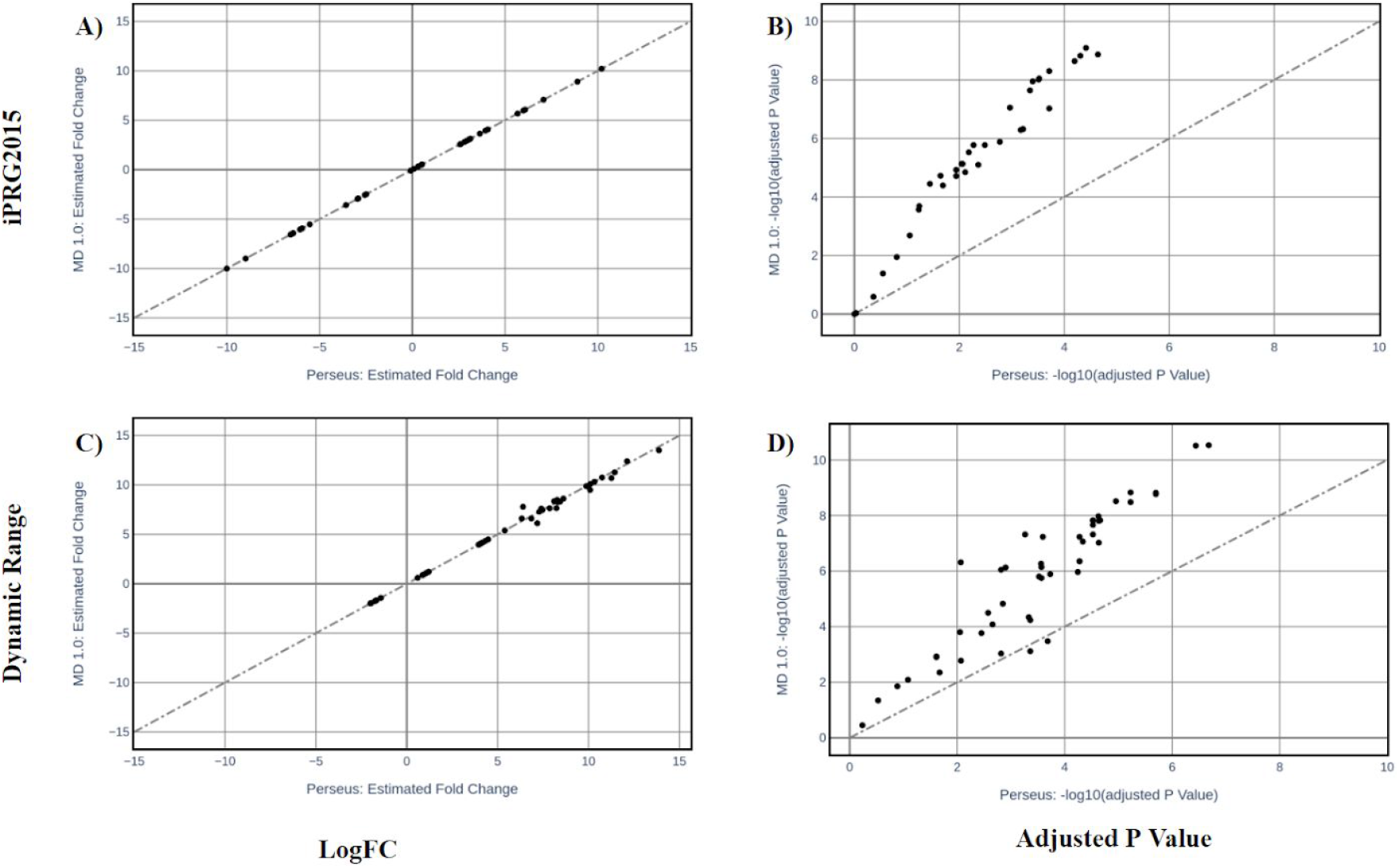
Comparison Analysis between LFQ results produced by Perseus and MD+ Discovery MD 1.0. Using the dynamic range dataset. (A) Scatter plot showing Log2 Fold Change Estimates (Pearson correlation = 0.998). (B) 2D Density Scatter plot showing Log10 Adjusted p-value estimates (Pearson correlation = 0.925). Using the iPRG2015 dataset (C) Scatter plot showing Log2 Fold Change Estimates (Pearson correlation = 1.000). (D) 2D Density Scatter plot showing Log10 Adjusted p-value estimates (Pearson correlation = 0.98).

**Figure 6.**
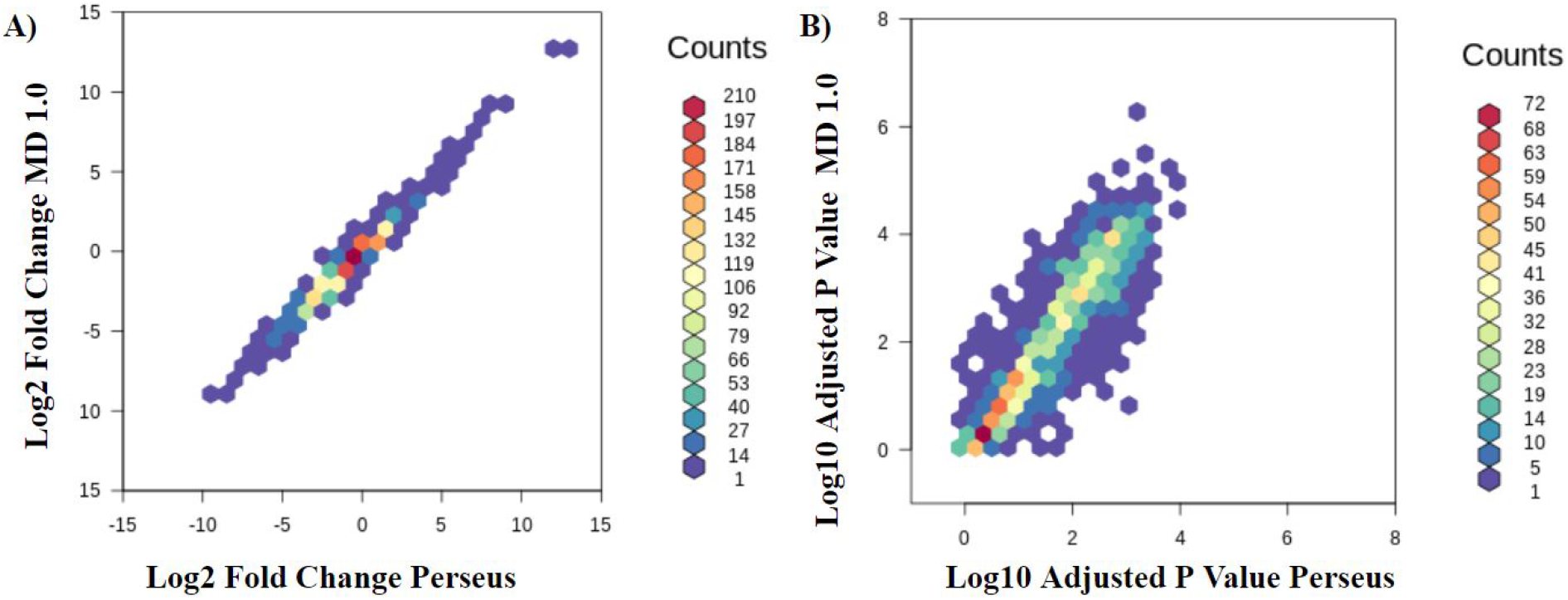
Comparison Analysis between LFQ results produced by Perseus and MD 1.0 using the HER dataset. (A) 2D Density Scatter plot showing Log2 Fold Change Estimates (Pearson correlation = 0.99). (B) 2D Density Scatter plot showing Log10 Adjusted p-value estimates (Pearson correlation = 0.90).

## Conclusion and Outlook

Mass Dynamics 1.0 is a web-based, integrated and automated analysis and collaboration environment that facilitates label free quantitative experiments. It currently accepts the MaxQuant output and is designed to be adaptable to accept data from other preprocessing tools. The output of MD 1.0 is served to the user via web pages which contain interactive figures, tables and downloads. The analysis performed by MD 1.0 is inherently reproducible both in platform and elsewhere, and can be easily shared or published to the community.

With the intention of broadening access to a wider range of users from computational and non-computational backgrounds, this environment provides many features that facilitate quality control, reproducibility, automation and transparency.

We demonstrated that MD 1.0 reproduces results similar to Perseus protocol reliably and accurately with respect to known benchmarking datasets, while producing comprehensive quality control reports that allows the user to be confident about the quality of experiment. MD 1.0 therefor constitutes a reliable, straight-forward and streamlined alternative to the Perseus platform when performing LFQ.

Mass Dynamics is well placed to begin expanding MD 1.0 in terms of analysis available beyond LFQ. Enrichment analysis such as Over-Representation analysis (ORA), Gene Set Enrichment Analysis (GSEA) might be achieved via integrations 3rd party databases such as GO, DAVID, KEGG or Drugbank. User interface improvements may involve more opportunities for annotation and sharing utilities. Lastly, leveraging the cloud-storage element of MD 1.0 implementation, insights may be gained by cross referencing experiment data and insights.

## Acknowledgements

The authors would like to thank members of the Future Industries Institute at the University of South Australia for their feedback and support during the development of MD+ Discovery. The authors would also like to thank the proteomics community for their ongoing development of comprehensive tools and repositories for the analysis and sharing of proteomics data without the work in this manuscript would be impossible.

## Author Contributions

Giuseppe Infusini wrote the MD 1.0 processing pipeline. Aaron Triantafyllidis built the web interface and set up the Amazon Web Services (AWS) compute and cloud storage infrastructure. Joseph Bloom performed the benchmarking analysis, prepared the figures and wrote the manuscript. Paula Burton, Giuseppe Infusini and Andrew Webb reviewed the manuscript. Paula Burton and Aaron Triantafyllidis interviewed and researched existing work in this space.

## Supplementary Figures and Tables

**Table S1:**
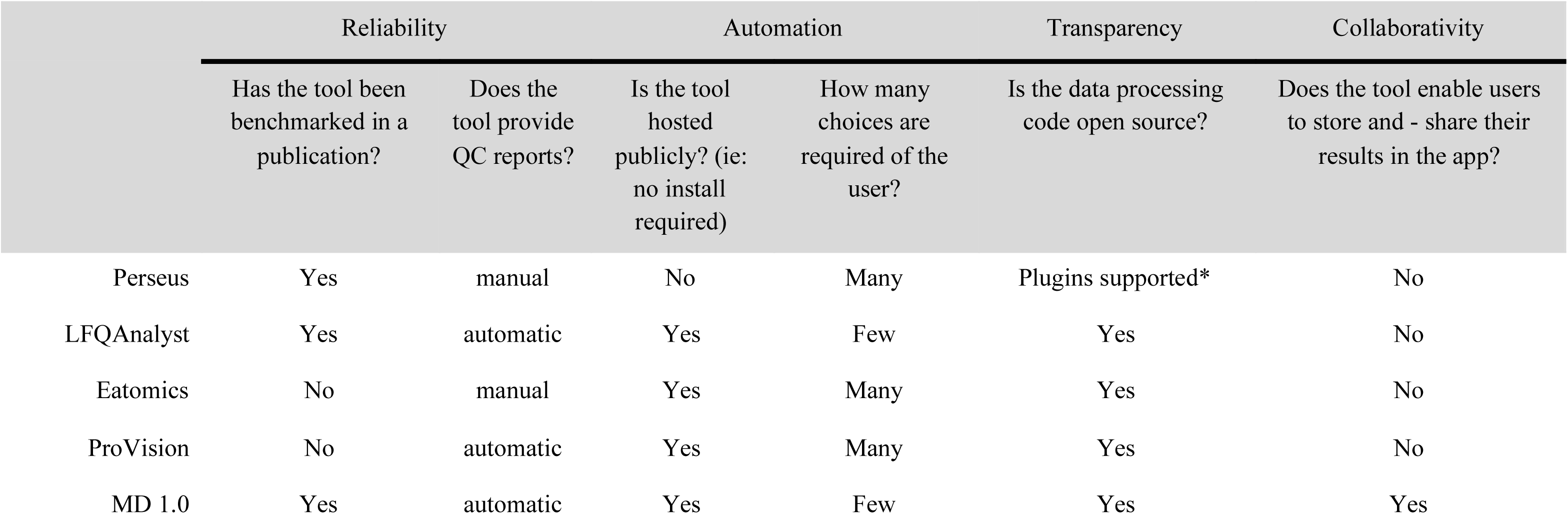
Qualitative Comparison of LFQ Statistics and Visualization Resources. *While we were not able to locate the source code for the Perseus computational platform, it is built using a plugin structure that users are able to contribute towards by building their own plugins in the C# programming language and more recently in PerseusNet the Python or R programming languages^7^.

|  | Reliability |  | Automation |  | Transparency | Collaborativity |
| --- | --- | --- | --- | --- | --- | --- |
|  | Has the tool been benchmarked in a publication? | Does the tool provide QC reports? | Is the tool hosted publicly? (ie: no install required) | How many choices are required of the user? | Is the data processing code open source? | Does the tool enable users to store and - share their results in the app? |
| Perseus | Yes | manual | No | Many | Plugins supported* | No |
| LFQAnalyst | Yes | automatic | Yes | Few | Yes | No |
| Eatomics | No | manual | Yes | Many | Yes | No |
| ProVision | No | automatic | Yes | Many | Yes | No |
| MD 1.0 | Yes | automatic | Yes | Few | Yes | Yes |

## References

1. Proteomics Technologies and Applications. (2019) doi:10.5772/intechopen.73445.

2. Wilcken, B., Wiley, V., Hammond, J. & Carpenter, K. Screening newborns for inborn errors of metabolism by tandem mass spectrometry. N. Engl. J. Med. 348, (2003).

3. Cox, J. et al. Accurate proteome-wide label-free quantification by delayed normalization and maximal peptide ratio extraction, termed MaxLFQ. Mol. Cell. Proteomics 13, 2513–2526 (2014).

4. Al Shweiki, M. R. et al. Assessment of Label-Free Quantification in Discovery Proteomics and Impact of Technological Factors and Natural Variability of Protein Abundance. J. Proteome Res. 16, 1410–1424 (2017).

5. Tyanova, S., Temu, T. & Cox, J. The MaxQuant computational platform for mass spectrometry-based shotgun proteomics. Nat. Protoc. 11, 2301–2319 (2016).

6. Tyanova, S. et al. The Perseus computational platform for comprehensive analysis of (prote)omics data. Nat. Methods 13, 731–740 (2016).

7. Rudolph, J. D. & Cox, J. A Network Module for the Perseus Software for Computational Proteomics Facilitates Proteome Interaction Graph Analysis. J. Proteome Res. 18, 2052–2064 (2019).

8. Shah, A. D., Goode, R. J. A., Huang, C., Powell, D. R. & Schittenhelm, R. B. LFQ-Analyst: An Easy-To-Use Interactive Web Platform To Analyze and Visualize Label-Free Proteomics Data Preprocessed with MaxQuant. J. Proteome Res. 19, 204–211 (2020).

9. Kraus, M., Mathew Stephen, M. & Schapranow, M.-P. Eatomics: Shiny Exploration of Quantitative Proteomics Data. J. Proteome Res. 20, 1070–1078 (2021).

10. Gallant, J. L., Heunis, T., Sampson, S. L. & Bitter, W. ProVision: a web-based platform for rapid analysis of proteomics data processed by MaxQuant. Bioinformatics 36, 4965–4967 (2020).

11. Smith, R. Conversations with 100 Scientists in the Field Reveal a Bifurcated Perception of the State of Mass Spectrometry Software. J. Proteome Res. 17, 1335–1339 (2018).

12. Quadroni, M. & James, P. Proteomics and automation. Electrophoresis 20, 664–677 (1999).

13. Pfeuffer, J. et al. OpenMS - A platform for reproducible analysis of mass spectrometry data. J. Biotechnol. 261, 142–148 (2017).

14. Creedon, H. et al. Identification of novel pathways linking epithelial-to-mesenchymal transition with resistance to HER2-targeted therapy. Oncotarget 7, 11539–11552 (2016).

15. Choi, M. et al. ABRF Proteome Informatics Research Group (iPRG) 2015 Study: Detection of Differentially Abundant Proteins in Label-Free Quantitative LC-MS/MS Experiments. J. Proteome Res. 16, 945–957 (2017).

16. The UniProt Consortium & The UniProt Consortium. UniProt: the universal protein knowledgebase. Nucleic Acids Research vol. 45 D158–D169 (2017).

17. Reactome - a curated knowledgebase of biological pathways. doi:10.3180/50.

